# Daily torpor reduces the energetic consequences of habitat selection for a widespread bat

**DOI:** 10.1101/2021.03.06.434212

**Authors:** Jesse M. Alston, Michael E. Dillon, Douglas A. Keinath, Ian M. Abernethy, Jacob R. Goheen

## Abstract

Homeothermy requires increased metabolic rates as temperatures decline below the thermoneutral zone, so homeotherms typically select microhabitats within or near their thermoneutral zones during periods of inactivity. However, many mammals and birds are heterotherms that relax internal controls on body temperature when maintaining a high, stable body temperature is energetically costly. Such heterotherms should be less tied to microhabitats near their thermoneutral zones, and because heterotherms spend more time in torpor and expend less energy at colder temperatures, heterotherms may even select microhabitats in which temperatures are well below their thermoneutral zones. We studied how temperature and daily torpor influence selection of diurnal roosts by a heterothermic bat (*Myotis thysanodes*). We (1) quantified the relationship between ambient temperature and daily duration of torpor, (2) simulated daily energy expenditure over a range of microhabitat (roost) temperatures, and (3) quantified the influence of roost temperature on roost selection. While warm roosts substantially reduced energy expenditure of simulated homeothermic bats, heterothermic bats modulated their use of torpor to maintain a constant level of energy expenditure over the course of a day. Daily torpor expanded the range of energetically economical microhabitats, such that roost selection was independent of roost temperature. Our work adds to a growing literature documenting functions of torpor beyond its historical conceptualization as a last-resort measure to save energy during extended or acute energetic stress.

## Introduction

The thermal environments in which organisms live strongly influence metabolic rates (Huey and Stevenson 1979, Brown et al. 2004, Pörtner and Knust 2007). Among homeotherms—which regulate body temperature internally within a narrow range to optimize physiological processes—metabolic heat production is tightly regulated in response to variation in temperature in the surrounding environment (i.e., ambient temperature; Lowell and Spiegelman 2000). Controlling body temperature thus requires increased energy expenditure by homeotherms when ambient temperatures depart from the thermoneutral zone (i.e., the range of ambient temperatures in which homeotherms can regulate body temperature with minimal metabolic effort; McNab 2002). Because survival and reproduction require that energy intake equal or exceed energy expenditure, operating in ambient temperatures outside the thermoneutral zone can reduce fitness over time (Angilletta et al. 2010, Boyles et al. 2011).

Although the influence of ambient temperature on metabolism in homeotherms is understood relatively well, many animals are heterotherms that can temporarily or partially allow body temperature to track ambient temperature (Withers et al. 2016). Heterothermy is common among mammals and birds (Geiser, 2004; Geiser and Ruf, 1995; McKechnie and Mzilikazi, 2011; Ruf and Geiser, 2015) and can reduce energy expenditure during both hot and cold periods (Stawski and Geiser 2012, Boyles et al. 2016, Nowack et al. 2017, Reher and Dausmann 2021). As ambient temperatures depart the thermoneutral zone, heterotherms can relax internal controls on metabolism; this physiological response allows body temperature to track ambient temperature and reduce or altogether eliminate the energetic costs of maintaining stable body temperatures outside the thermoneutral zone (Levesque et al. 2016). Heterotherms often achieve this by entering daily torpor, a short-term hypometabolic state of inactivity (Ruf and Geiser 2015).

Heterotherms use daily torpor more as ambient temperatures decline below the thermoneutral zone (Chruszcz and Barclay, 2002; Geiser and Broome, 1993; Rambaldini and Brigham, 2008; Solick and Barclay, 2006), but it is unclear how this tendency translates to differences in energy expenditure across differences in temperature. For a given period of time, total energy expenditure for heterotherms depends on (1) the duration and frequency of bouts of torpor, (2) ambient temperatures, and (3) the difference in metabolic rates between torpor and homeothermy at a given ambient temperature. Energy expenditure might increase as ambient temperatures fall below the thermoneutral zone: even though heterotherms save energy by using torpor, declines in energy expenditure from using torpor more when it is cold do not fully compensate for the increased energetic costs of maintaining homeothermy in colder ambient temperatures (Fig. 1B). In this scenario, periodic bouts of torpor dampen but do not completely offset increases in energy expenditure during periods of homeothermy at cold ambient temperatures. Alternatively, it is possible that energy expenditure by heterotherms is stable through a wide range of ambient temperatures because energy savings from using progressively more torpor at progressively colder ambient temperatures closely matches increases in energy expenditure from maintaining homeothermy at colder ambient temperatures (Fig. 1C). Finally, as ambient temperatures decline, the energetic savings from torpor could more than offset the increased energy expenditure necessary to maintain homeothermy (Fig. 1D).

**Fig. 1.**
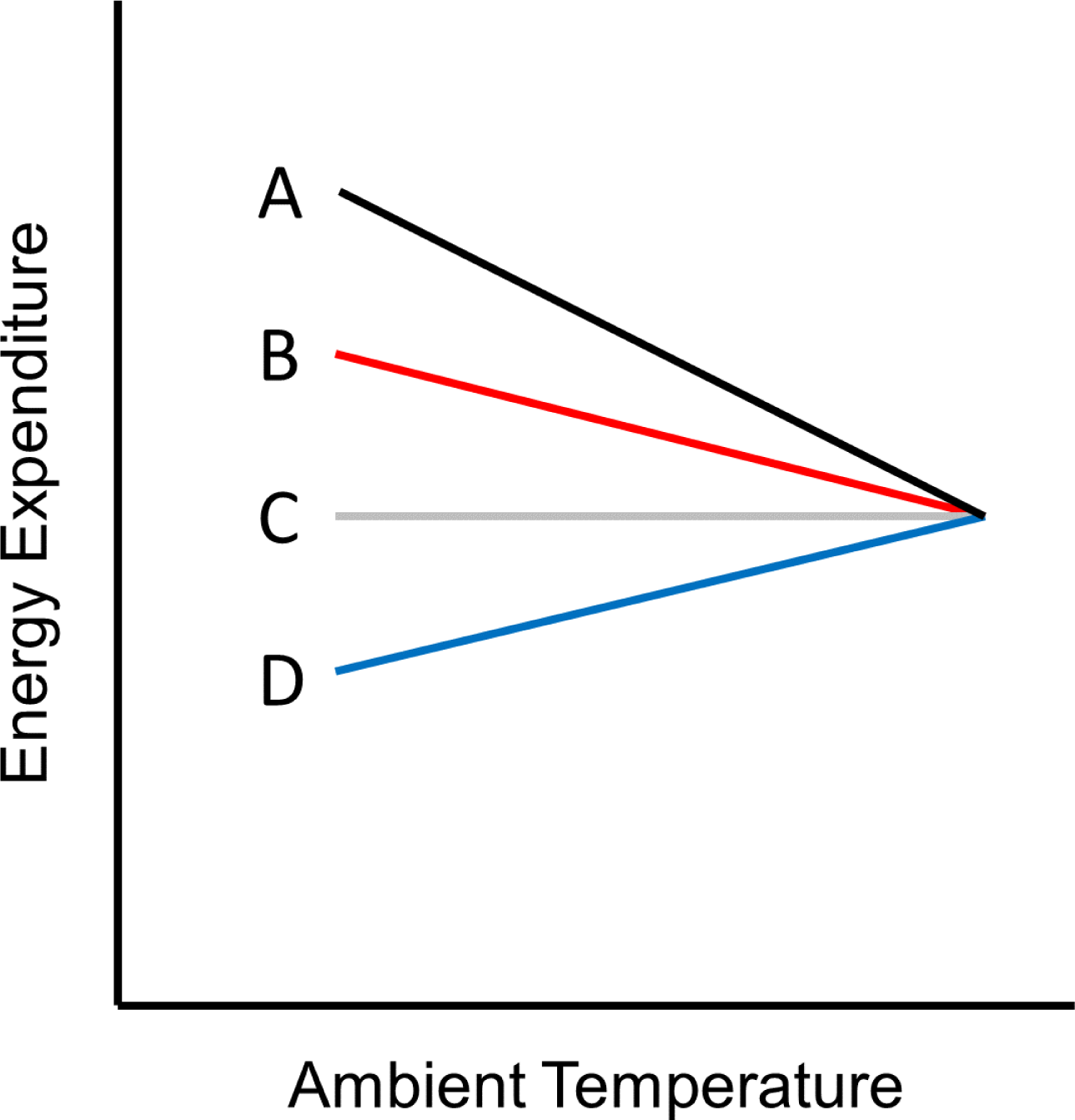
Heuristic diagram outlining the potential energetic benefit to an individual bat of using periodic bouts of daily torpor rather than remaining in homeothermy at ambient temperatures below the thermoneutral zone. This diagram is similar to a classic Scholander curve except for one detail: while a Scholander curve illustrates metabolic rate or energy expenditure at a constant ambient temperature and physiological state (i.e., either homeothermy or torpor) in laboratory conditions, this diagram illustrates energy expenditure when ambient temperature and physiological state vary through time as they do in field conditions. Specifically, this diagram assumes that (1) bats use more torpor when it is cold than when it is warm, (2) ambient temperatures vary over the course of the day, and (3) ambient temperatures below the thermoneutral zone are more prevalent than ambient temperatures above the thermoneutral zone (see Cunningham et al. 2021). Each hypothetical relationship would result in a different pattern of roost selection by animals seeking to minimize energy expenditure during periods of inactivity. The black (A) line represents energy expenditure over a day while maintaining homeothermy 100% of the time (i.e., never using torpor). The red (B), grey (C), and blue (D) lines indicate energy expenditure over a day while using some amount of torpor. For all three relationships, torpor provides energy savings (i.e., the difference between the black and other lines), and these savings are most pronounced at colder ambient temperatures. (B) For bats that use torpor, energy expenditure *increases* at colder ambient temperatures because while some energy is saved from employing torpor, maintaining homeothermy is more costly at colder than at warmer ambient temperatures. A bat exhibiting this relationship should select *warm* roosts to reduce energy use. (C) For bats that use torpor, energy expenditure *is stable* across a wide range of ambient temperatures because the energy saved from employing torpor matches (and thus offsets) the increase in energy expended to maintain homeothermy at colder temperatures. A bat exhibiting this relationship should not benefit from selecting either warm or cool roosts, and should thus select neither warm nor cool roosts. (D) For bats that use torpor, energy expenditure *decreases* at colder ambient temperatures because relatively more energy is saved from using torpor even as maintaining homeothermy is more costly at colder than at warmer ambient temperatures. A bat exhibiting this relationship should select *cool* roosts to reduce energy use.

Such relationships between ambient temperature and energy expenditure have cascading repercussions for other aspects of an animal’s life. For example, ambient temperature often influences habitat selection by animals seeking to minimize energy expenditure (e.g., Huey 1991, Freitas et al. 2016, Sarmento et al. 2019, Alston et al. 2020). Homeotherms have relatively fixed relationships between ambient temperature and metabolic rate, and thus often consistently select habitats to maintain optimal body temperatures with little metabolic effort (e.g., Poole et al. 2016, Courbin et al. 2017, Sarmento et al. 2019). In contrast, looser relationships between ambient temperature and metabolic rate for heterotherms may allow heterotherms to select habitats with less regard to ambient temperature, or even to prefer habitats that might be colder than optimal for homeotherms. For example, heterothermic Australian owlet-nightjars (*Aegotheles cristatus*) preferentially roost in colder, less thermally stable tree cavities, whereas homeothermic cavity-nesting birds typically select warmer, more thermally stable tree cavities (Doucette et al. 2011). Empirical data on habitat selection by heterotherms is rare, however, particularly for free-ranging animals.

Uncertainty surrounding the form and strength of relationships between ambient temperature and energy expenditure limit our understanding of temperature-driven habitat selection by heterotherms. For an animal attempting to minimize energy expenditure during periods of inactivity, each of the hypothetical relationships between energy expenditure and ambient temperature in Fig. 1 would result in a different pattern of habitat selection. A heterotherm exhibiting the relationship shown by the red (B) line in Fig. 1 should select warm microhabitats to save energy, similar to homeotherms. A heterotherm exhibiting the relationship shown by the grey (C) line in Fig. 1 should not select microhabitats based on their thermal characteristics. This pattern of habitat selection would also diverge from the pattern followed by homeotherms. A heterotherm exhibiting the relationship shown by the blue (D) line in Fig. 1 should select cool microhabitats to save energy, opposite of the pattern followed by homeotherms. Empirical tests of the influence of ambient temperature on energy expenditure are thus needed to understand how ambient temperature drives habitat selection for heterotherms.

We sought to understand how ambient temperature influences energy expenditure, and how energy expenditure in turn influences habitat selection, in a bat that is widely distributed throughout western North America (fringed myotis, *Myotis thysanodes*). Like other bats inhabiting temperate latitudes, fringed myotis are heterotherms that are believed to select diurnal roosts to minimize energy expenditure during diurnal periods of inactivity (Sedgeley 2001, Willis and Brigham 2005, Ruczyński 2006). At temperate latitudes, temperature within roosts can vary substantially throughout the day and year, and ambient temperature influences the amount of time bats spend in torpor each day. Like other heterotherms, bats spend more time in torpor when it is cold than when it is hot (Chruszcz and Barclay 2002, Solick and Barclay 2006, Rambaldini and Brigham 2008). We hypothesized that differences in energy expenditure at roosts of varying temperatures drive patterns of roost selection (i.e., bats select roosts that minimize energy expenditure). Specifically, we weighed evidence for four competing sets of predictions (Fig. 2):

**Fig. 2.**
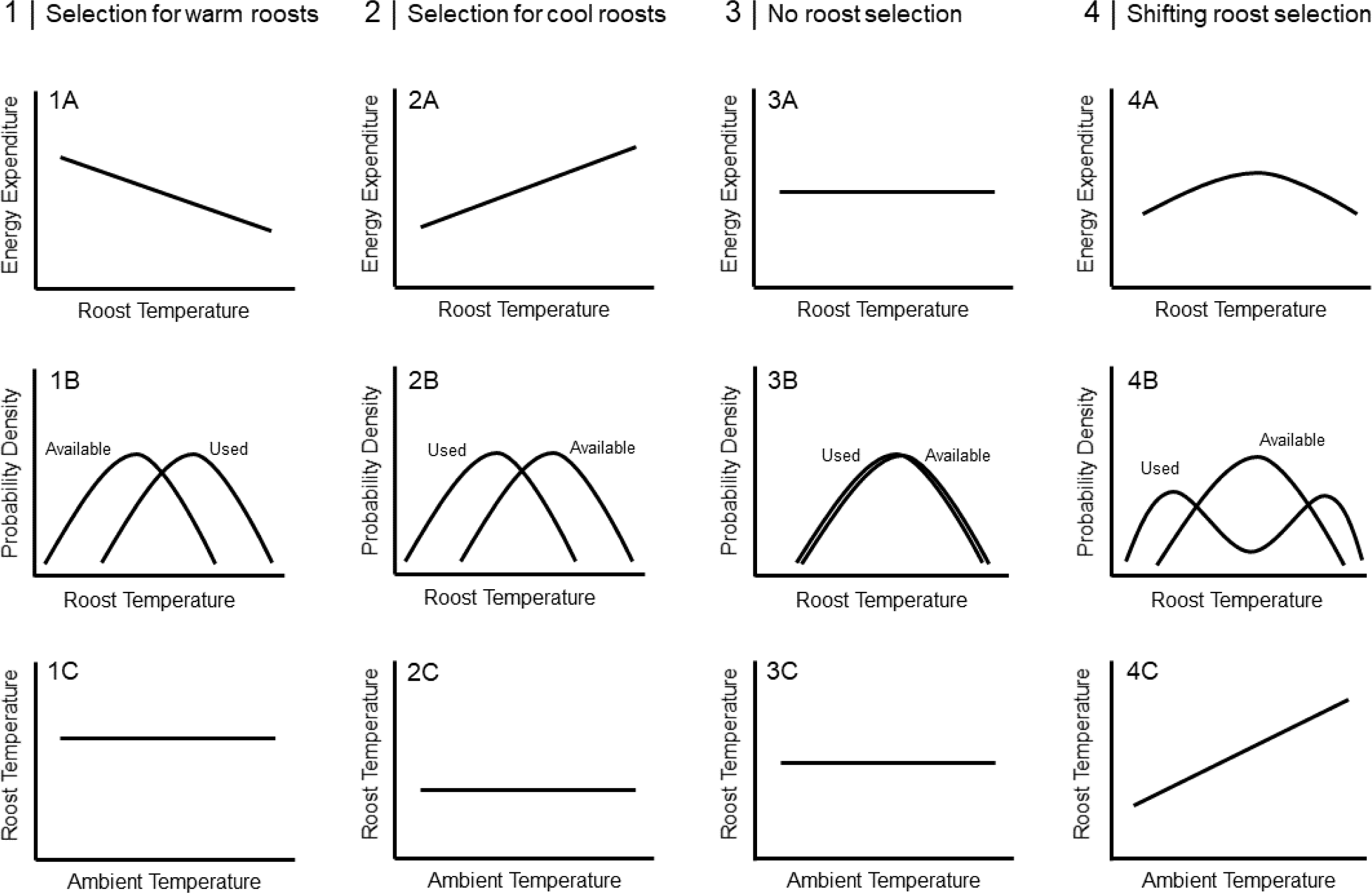
Four competing sets of predictions of roost selection by a heterothermic bat. Each column represents one of four sets of predictions, and each row represents a statistical relationship consistent with the predictions. In column 1, energy expenditure over the course of a day is higher in cool roosts than in warm roosts (1A). In response, bats select warm roosts to minimize energy expenditure during the day (1B). In this scenario, there should be no directional relationship between ambient temperature and roost temperature (i.e., bats always select warm roosts regardless of ambient temperature; 1C). In column 2, energy expenditure over the course of a day is higher in warm roosts than in cool roosts (2A). In response, bats select cool roosts to minimize energy expenditure during the day (2B). In this scenario, there should be no directional relationship between ambient temperature and roost temperature (i.e., bats always select cool roosts regardless of ambient temperature; 2C). In column 3, energy expenditure over the course of a day is constant across roosts of all temperatures (because bats can adaptively use torpor so that roost temperatures over the course of a day have little influence on overall energy expenditure; 3A). Because energy expenditure is consistent across roosts of all temperatures, bats do not select roosts due to roost temperature (3B). In this scenario, there is no relationship between ambient temperature and roost temperature (i.e., bats never select roosts due to temperatures within roosts, regardless of ambient temperature; 3C). In column 4, energy expenditure peaks at intermediate roost temperatures where bats use relatively little torpor but the costs of maintaining homeothermy are relatively high (4A). In response, bats select cool roosts on cool days and warm roosts on warm days (4B) because torpor saves more energy in cool roosts than in warm roosts. In this scenario, the relationship between ambient temperature and roost temperature should be positive (i.e., bats select warmer roosts on warmer days; 4C).

*Prediction Set 1*: Bats select warm roosts regardless of ambient temperature. In this scenario, energy expenditure during the day should be higher in cool roosts than in warm roosts (Fig. 2.1A) because the energetic benefits from being warmer when bats are maintaining homeothermy outweigh the energetic costs of spending less time in torpor. If this is the case, bats should select roosts that are warmer compared to available structures on the landscape (Fig. 2.1B); this pattern of selection should be consistent regardless of ambient temperature during the day (Fig. 2.1C).

*Prediction Set 2*: Bats select cool roosts regardless of ambient temperature. In this scenario, energy expenditure during the day should be higher in warm roosts than in cool roosts (Fig. 2.2A) because the energetic benefits from spending more time in torpor outweigh the energetic costs of being colder when bats are maintaining homeothermy. If this is the case, bats should select roosts that are cooler compared to available structures on the landscape (Fig. 2.2B); this pattern of selection should be consistent regardless of ambient temperature during the day (Fig. 2.2C).

*Prediction Set 3*: Bats do not alter roost selection as ambient temperatures change. In this scenario, energy expenditure during the day is roughly equal across roosts of all temperatures (Fig. 2.3A). This could occur if bats modulate use of torpor such that roost temperatures over the course of a day have little influence on overall energy expenditure. In this case, bats should select roosts that are similar in temperature to available structures on the landscape (Fig. 2.3B), and this pattern of selection should be consistent regardless of ambient temperature during the day (Fig. 2.3C).

*Prediction Set 4*: Bats select cool roosts on cool days and warm roosts on warm days (shifting roost selection). In this scenario, energy expenditure is lower in cool roosts than in warm roosts on cool days, lower in warm roosts than in cool roosts on warm days, and consistently higher in roosts at intermediate ambient temperatures (Fig. 2.4A). This may arise because of threshold effects from a non-linear relationship between ambient temperature and torpor use. Namely, a threshold may exist above which homeothermy requires relatively little energy even as bats spend little time in torpor, but below which bats save a substantial amount of energy by using torpor. Near the threshold, however, bats may use relatively little torpor even as maintaining homeothermy is relatively energetically costly. In this case, bats should select roosts that are roughly the same temperature on average as available structures on the landscape (though the distribution may be bimodal; Fig. 2.4B), and temperatures in roosts should be positively correlated with ambient temperature (Fig. 2.4C).

## Methods

### Study Area and Species

We conducted our study during the summers of 2017 and 2018 on Jewel Cave National Monument (43° 45’ N, 103° 45’ W) and surrounding areas of Black Hills National Forest in South Dakota, USA. Our study area is described in Alston et al. (2019). Mean monthly summer high temperatures range between 22 – 27°C and mean monthly summer precipitation ranges between 60 – 80 mm (Western Regional Climate Center 2018). Open ponderosa pine (*Pinus ponderosa*) forests dominate, with Rocky Mountain juniper (*Juniperus scopulorum*) and quaking aspen (*Populus tremuloides*) occurring locally. Forests are actively managed to prevent wildfire, and those managed by the US Forest Service and private landowners also undergo intensive logging. Forests form a mosaic with northern mixed-grass prairie where a large stand-replacing fire occurred in in 2000. A large network of caves lie underground, and the landscape exhibits substantial topographic relief in the form of intersecting canyon systems and rock outcrops.

Fringed myotis roost in caves, mines, rock crevices, tree cavities, and under the sloughing bark of dead trees, and forage in forest canopy and riparian areas (O’Farrell and Studier 1980). We chose to focus on males because sex ratios of bats in the Black Hills are heavily (>90%) male-biased (a common pattern in high-elevation areas; Barclay, 1991; Cryan et al. 2000; Senior et al. 2005), because male *M. thysanodes* usually roost solitarily (O’Farrell and Studier 1980), and because male bats maintain consistent patterns of torpor use throughout the reproductive season (unlike females, which alter patterns of torpor use at different stages of reproduction; Chruszcz and Barclay, 2002; Dzal and Brigham, 2013; Johnson and Lacki, 2014).

### Capture and VHF Telemetry

We used mist nets to capture bats over permanent and semi-permanent water sources (e.g., springs, stock tanks, and stock ponds). From June through August of 2017 and 2018, we netted 20 and 49 nights, respectively, at 15 water sources. We opened mist nets at civil sunset and closed them after five hours or during inclement weather.

We affixed temperature-sensitive VHF transmitters (LB-2XT model .28/.33 g – Holohil Systems Ltd., Carp, ON, Canada) between the scapulae of adult male fringed myotis with latex surgical adhesive (Osto-Bond, Montreal Ostomy, Montreal, QC, Canada). The transmitters measure and transmit data on skin temperature—an accurate proxy for body temperature—of bats, enabling researchers to delineate bouts of torpor (Barclay et al. 1996, Chruszcz and Barclay 2002, Stawski and Geiser 2010). All transmitters weighed <5% of the mass of the bat (Aldridge and Brigham 1988). We tracked bats to roosts each day transmitters were active, and installed VHF data loggers (SRX800-D1 – Lotek Wireless Inc., Newmarket, ON, Canada) that collected and recorded data transmitted by the VHF transmitters. All protocols were approved by the University of Wyoming and National Park Service Animal Care and Use Committees and met guidelines approved by the American Society of Mammalogists for research on wild mammals (Sikes and the Animal Care and Use Committee of the American Society of Mammalogists 2016).

### Energetic Modelling

To quantify torpor use, we delineated bouts of torpor from data logger readings that captured full days (i.e., from roost entry in the morning to roost exit in the evening) of skin temperature data from individual bats. This was a fraction of total days in which we located roosts, because bats typically were not located until after they entered roosts. We defined torpor as beginning when skin temperature dropped below the lowest skin temperature of bats maintaining homeothermy during a day and ending when skin temperature began a steep rise that led to bats re-entering homeothermy or leaving a roost (as recommended by Barclay et al. 2001; Appendix S1: Fig. S1). Because fat reserves and body mass can substantially alter the amount of time spent in torpor (Wojciechowski et al. 2007, Stawski and Geiser 2010, Vuarin et al. 2013), we also controlled for the body mass of each individual at time of capture on torpor duration. We then used the modelling software ‘Stan’ (Carpenter et al. 2017) via the R package ‘brms’ (v2.13.0; Bürkner 2017) to build a linear Bayesian hierarchical model to quantify the influence of ambient temperature and body mass on torpor duration while accounting for non-independence among data points collected from the same individual. The model included 3 chains run for 13,000 iterations (1,000 iterations of warm-up and 12,000 iterations of sampling). We assessed chain convergence using the Gelman-Rubin diagnostic (Ȓ) and precision of parameter estimation using effective sample size. Ȓ < 1.01 and effective sample sizes > 10,000 represent acceptable convergence and parameter precision (Gelman et al. 2013, Kruschke 2015). We used leave-one-out cross validation to check model fit using the R packages ‘loo’ (v2.2.0; Vehtari et al. 2017) and ‘bayesplot’ (v1.7.2; Gabry et al. 2019) to visually assess the cross-validated probability integral transform.

To quantify energy expenditure in bats, we combined published estimates of metabolic rates of fringed myotis as a function of temperature (Studier and O’Farrell 1976) and the linear model of the influence of ambient temperature on torpor use to simulate the influence of roost temperature on energy expenditure. Specifically, we simulated minute-by-minute energy expenditure by bats in each used roost between 0445 hrs and 2100 hrs (typical entry and exit times for bats in our study) on each day over the duration of our study period. We modeled torpor use as a function of decision rules that reflect torpor use observed over the course of our study (raw data presented in Appendix S1: Table S1). Specifically, we assumed that bats entered torpor immediately upon entering roosts, exited torpor after an interval determined by roost temperature, and remained in homeothermy for the rest of the time spent in the roost except for a shorter bout of torpor in the evening. We further assumed that bats would use 86.9% of the duration of daily torpor in the morning and 13.1% in the afternoon unless the afternoon bout of torpor would be less than 30 minutes in duration, in which case 100% of the day’s torpor would occur in the morning period. We also assumed that the mean duration of torpor that we observed would be used in the baseline “average” roost, with the duration of torpor in warmer and cooler roosts determined by the slope of the modeled relationship between ambient temperature and torpor use described in the above paragraph. To account for uncertainty in our estimate of the slope of the relationship between ambient temperature and daily torpor use, for each roost on each day we randomly drew a different slope estimate for this relationship from the posterior distribution of slope estimates from the model described in the prior paragraph.

### Roost Characterization

To characterize rock roost structures, we collected data for 31 roosts and 62 randomly sampled available (i.e., unused by bats in our study) roosts. Hereafter, we distinguish between ‘used roosts’ and available but unused ‘available roosts’; we use the term ‘roost structure’ when we refer to both used and available roosts simultaneously. We identified available rock roosts in two ways: at each used roost, we 1) located the nearest rock crevice large enough to hold a bat, and 2) generated a paired point in a random cardinal direction a random distance between 100 – 300 m away, then located the nearest rock crevice large enough to hold a bat.

To characterize tree roost structures, we collected data for 9 used roosts and 36 randomly sampled available roosts. We identified available tree roosts in two ways: at each used roost, we 1) located the nearest snag and selected the nearest cavity large enough to hold a bat, and 2) generated a paired point in a randomly determined distance between 100 – 300 m away, in a randomly-determined (cardinal) direction, then located the nearest tree cavity large enough to hold a bat. For each available point, we placed data loggers in two locations: one in a cavity in the trunk and one underneath sloughing bark. We defined available roost trees as any dead tree with a visible defect (e.g., sloughing bark or cavities) sufficiently large to hold a bat. This description fit every tree in which we found a bat roosting.

In Summer 2018, we monitored temperatures within both used and available roosts using data loggers (Model MX2201; Onset Computer Corporation, Bourne, MA, USA). The first data loggers were deployed on 17 July 2018, and the last data logger was removed on 8 October 2018. This period of time includes the full range of daily high temperatures occurring during the active season for bats at our study site. During data logger deployment and opportunistically thereafter, we checked roost structures for the presence of bats. We sometimes found bats in used roosts, but we never found bats in available roosts. When we found bats in used roosts, we waited to deploy data loggers until there was no bat within the roost.

To quantify the thermal characteristics of each roost structure, we calculated the mean temperature within each roost structure for periods between 0445 and 2100 hrs, which corresponds with the period in which a bat is likely to be within a roost (Appendix S1: Table S1). To control for potential confounding variables, we also calculated the timing of the peak temperature in all roost structures (because if two roost structures have the same mean temperature but peak in temperature at different times, the roost structure with the later peak will have cooler temperatures in the morning when bats use torpor most), and the standard deviation of temperature during the day (because stability in roost temperature can be an important factor in roost selection; Sedgeley, 2001). To quantify the timing of the daily temperature peak, we located the peak temperature in each roost structure for each day and calculated the mean time of day at which this occurred over our study period. To quantify thermal stability in roost structures, we calculated the standard deviation of temperatures between 0445 and 2100 hrs in each roost structure for each day and calculated the mean daily standard deviation over our study period. To ensure consistency, we only calculated these values for the period between July 28 and September 31 (a period in which all data loggers were actively logging temperatures, and in which average daily high temperatures correspond with the range a bat might be exposed to during the active season in our study area).

We used the R statistical software environment (R Core Team 2020) to quantify differences between used and available roosts. To determine whether bats select cooler roosts than those available, we used the modelling software ‘Stan’ (Carpenter et al. 2017) via the R package ‘brms’ (v2.13.0; Bürkner 2017) to build a binomial-family Bayesian model to quantify the influence of mean temperature within roost structures, the timing of daily peaks in temperature within roost structures, and the standard deviation of temperatures within roost structures on roost selection. The model included 3 chains run for 13,000 iterations (1,000 iterations of warm-up and 12,000 iterations of sampling). We assessed chain convergence using Ȓ and precision of parameter estimation using effective sample size. We checked predictive performance with receiver operating curve analysis using the R package ‘pROC’ (v1.16.2; Robin et al. 2011) and used the R package ‘bayesplot’ (v1.7.2; Gabry et al. 2019) to visually assess binned residual plots.

## Results

We tracked 46 bats to 107 roosts (93 in rock crevices and 14 in trees) and collected 27 full days of skin temperature data from 7 bats. Data from 16 data loggers within roost structures (3 used rock, 12 available rock, 1 available tree) could not be collected because they were not relocated or were dislodged from roost structures. We thus excluded these data from analyses, leaving a total of 122 (78 rock, 44 tree) data loggers that collected data on temperatures within roost structures.

Use of torpor stabilized daily energy expenditure across the range of roost temperatures observed during our telemetry study. In our model of the effect of ambient temperature on daily torpor duration, 95% credible intervals for the effect of mean ambient temperature over the course of the day on daily torpor duration did not cross 0 (parameter estimate: −37.4 min; 95% credible intervals: −64.0 – −12.6 min), indicating that bats spent ca. 37 minutes less in torpor per day for each additional 1°C in daily mean ambient temperature between 0445 hrs and 2100 hrs (Fig. S2). Assessment of the cross-validated probability integral transform indicated that model fit was adequate. When incorporated into our simulation of bat energy expenditure over the course of a typical day, this estimate of the relationship between ambient temperature and torpor use led to similar estimates of energy expenditure across temperatures within used roosts (Fig. 3; blue points). Daily energy expenditure was roughly equivalent in all roosts. Our estimates for energy expenditure using observed bat behavior were always substantially lower and less variable than our estimates for energy expenditure if bats had remained in homeothermy all day (Fig. 3; red points). Bats that remain in homeothermy would expend substantially more energy in cool roosts than warm roosts.

**Fig. 3.**
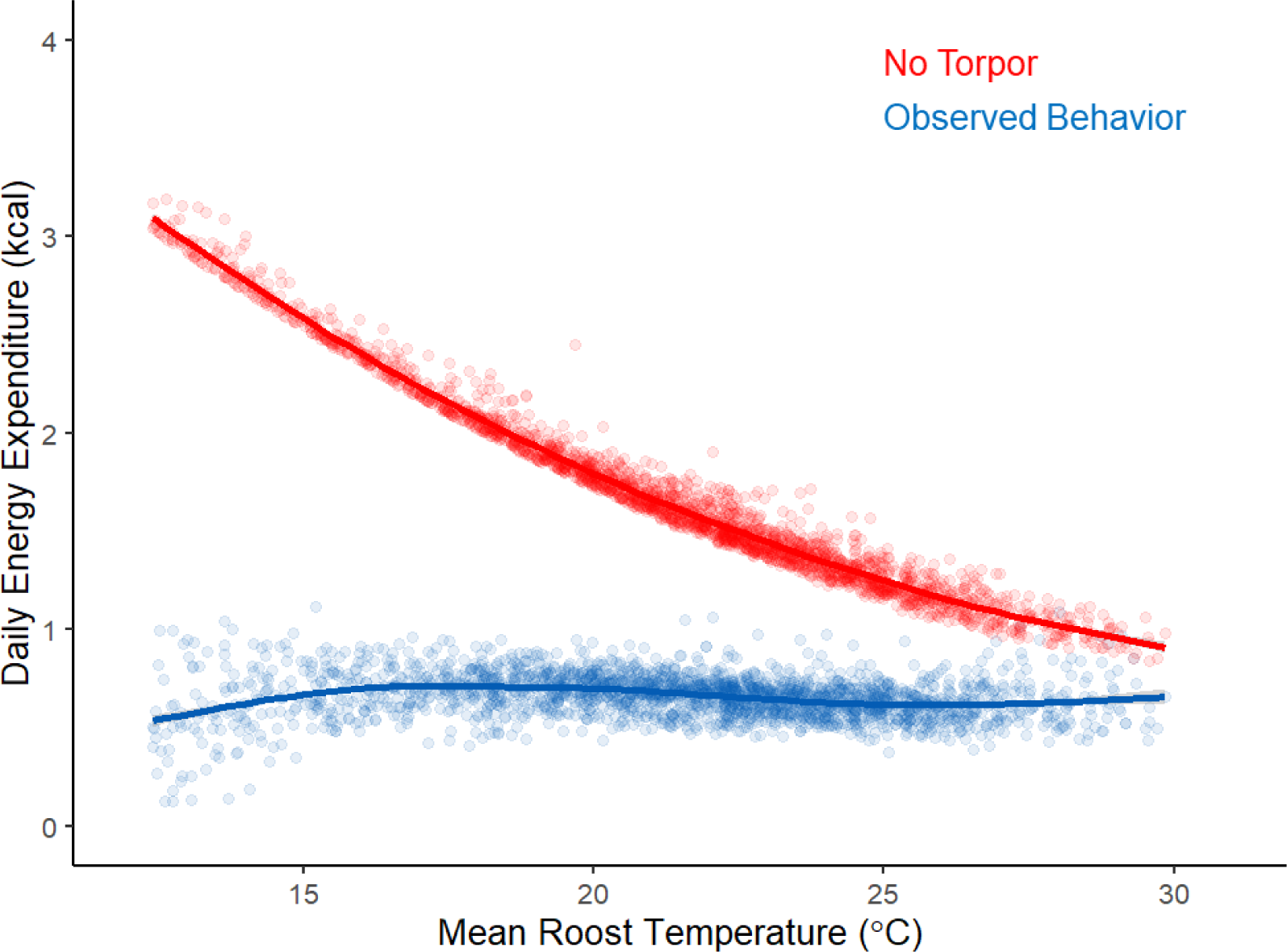
Results of our simulation of daily energy expenditure by fringed myotis over the range of temperatures observed in used roosts. Each point represents one day. The red points represent estimated daily energy expenditure if bats never used torpor. The blue points represent our estimate of energy expenditure over the course of a day if part of the day is spent in torpor (with daily duration of torpor a function of daily ambient temperature as observed in our study). The lines represent loess regressions of the relationship between roost temperature and daily energy expenditure. Estimates of daily energy expenditure incorporating observed bat behavior are steady across all roost temperatures observed during our study. The blue points in this figure correspond with Row A in Fig. 2, and are most closely matched by Fig 2.3A.

Overall, temperatures in both rock and tree roost structures were similar, though roost structures in trees were slightly cooler and less stable than roost structures in rocks. We therefore pooled rock and tree roost structures in roost selection analyses, but we report descriptive statistics for each type of roost structure in Appendix S1.

Despite substantial variation in temperatures among roost structures, we found little evidence that the thermal characteristics of used roosts differed from those of available roosts (Fig. 4). In our model of roost selection, 95% credible intervals for the effect of mean ambient temperature over the course of the day on roost selection did not cross 0 (parameter estimate: 0.30; 95% credible intervals: 0.04 – 0.58), indicating that bats were more likely to roost in warm roost structures than cool ones. However, predictive performance was poor (AUC: 0.650), and overall, used roosts (20.1°C) had similar mean temperatures as available roosts (19.4°C; Fig. 4A). Bats also did not differentiate between roost structures with temperatures peaking late in the day versus roost structures with temperatures peaking early in the day (Fig 4B). In our model of roost selection, 95% credible intervals for the effect of the timing of daily peaks in temperature on roost selection crossed 0 (parameter estimate: −0.10; 95% credible intervals: −0.34 – 0.14). Overall, used roosts (1408 hrs) peaked in temperature at similar times as available roosts (1434 hrs). Bats also did not differentiate between roosts with stable temperatures and those with more variable temperatures (Fig. 4C). In our model of roost selection, 95% credible intervals for the effect of standard deviation in roost temperature over the course of the day on roost selection crossed 0 (parameter estimate: −0.20; 95% credible intervals: −0.47 – 0.06) Overall, there was no difference in the standard deviation of temperatures of used roosts (7.0°C) and available roosts (7.0°C). Finally, there was also no relationship between ambient temperature on a given day and mean temperatures within roosts used on that day (*R*^2^ = 0.03; *p* = 0.132; Fig. 5).

**Fig. 4.**
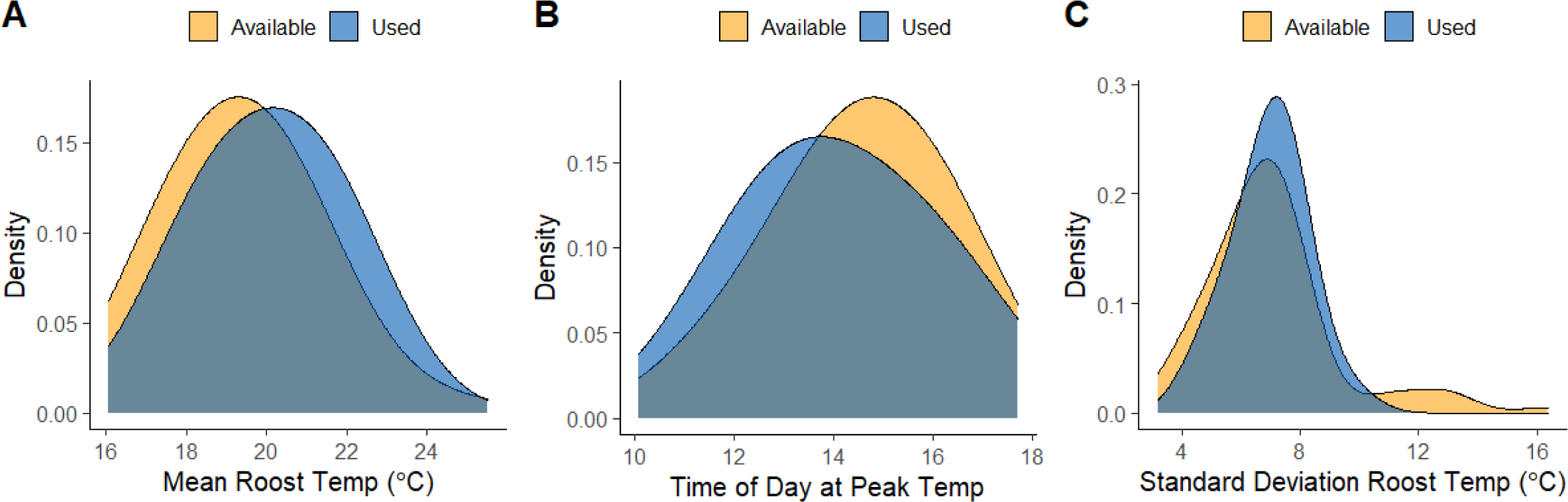
Kernel density plots comparing thermal characteristics within used and available roost structures: mean temperature (A), time of day at peak temperature (B), and the standard deviation of temperature (C). Blue distributions represent used roosts, while orange distributions represent available roosts. These plots illustrate the results of our binomial model of roost selection. Used roosts were slightly warmer on average than available roosts, but their distributions largely overlapped (A). Temperatures peaked slightly earlier in used roosts than available roosts, but this was a function of temperatures in warmer roosts tending to peak earlier in the day (*r* = −0.19 for the relationship between mean temperature within roost structures and time of day at peak temperature) and their distributions largely overlap (B). The standard deviation in temperatures within used roosts is very similar to the standard deviation in temperatures within available roosts, although bats did not use the few roost structures with very high standard deviations (C). Panel A in this figure corresponds with Row B in Fig. 2, and is most closely matched by Fig. 2.3B.

**Fig. 5.**
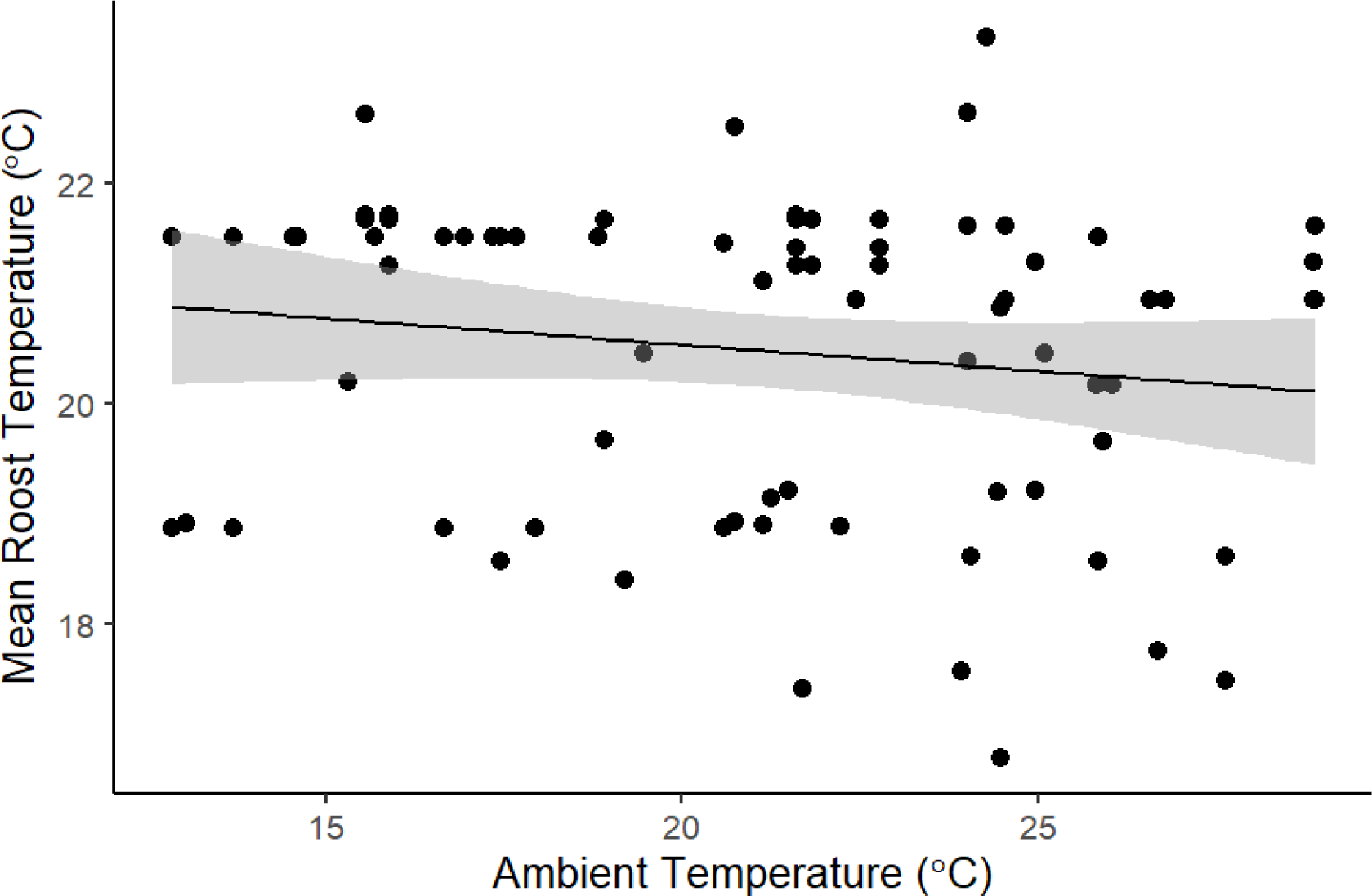
Scatter plot of the relationship between ambient temperature on a given day and mean temperature within used roosts. Each point is based on observed data, and represents a roost used for one day; some roosts (n = 14) were used on multiple days and are thus represented by multiple points on this plot. The line represents the regression line for this relationship and the grey band represents 95% confidence intervals. Ambient temperature on a given day did not influence whether bats used warm or cool roosts (*p* = 0.06; *R*^2^ = 0.04). This figure corresponds with Row C in Fig. 2, and is most closely matched by Fig. 2.3C.

## Discussion

The thermal environments in which animals operate strongly influence physiological processes, and can thereby pose substantial challenges in variable environments. How animals overcome these challenges is a central question in animal ecology. Attempts to address this question have focused largely on poikilotherms and homeotherms. Because heterotherms are neither as strongly tied to narrow ranges of body temperature as homeotherms nor as subject to ambient temperatures as poikilotherms, the relationships between temperature and habitat selection for heterotherms should differ fundamentally from those of either homeotherms or poikilotherms. Specifically, whereas homeotherms select microhabitats near the thermoneutral zone during periods of inactivity, heterotherms should have less incentive to do so.

We sought to better understand how variation in ambient temperature influences use of daily torpor and habitat selection for heterotherms, using a species of bat as a model system. Simulations of energy expenditure at varying roost temperatures revealed that bats can modulate use of torpor to maintain constant energy expenditure over the course of a day over a wide range of temperatures within roosts. As a result, roost selection was not driven by temperatures within roosts. Our results provide evidence for Prediction Set 3 (no selection) in our introduction (Fig. 2).

The energetic savings associated with torpor—particularly at cooler temperatures—likely result in habitat selection that differs substantially from habitat selection by homeotherms. For example, we showed that use of daily torpor can reduce the energetic costs of inhabiting roosts that are colder than optimal for homeotherms. If bats were strict homeotherms, the energetic costs of inhabiting cool roosts would have been substantially higher (Fig. 3), which would likely result in bats selecting warm roosts. In contrast, heterothermic bats face little pressure to select warm habitats, even on relatively cool days. Daily torpor does not simply loosen the thermal constraints facing homeotherms at temperatures below the thermoneutral zone—it can entirely mitigate them. Additional studies of the relationships between temperature, torpor use, and habitat selection would be valuable for establishing the generality of this finding for other heterothermic species.

Individual traits (e.g., sex, age, and reproductive condition) can alter the energetic costs and benefits of using torpor for heterotherms, thereby driving divergence from the pattern demonstrated in this study. For example, roost selection by bats varies by sex, age, and reproductive condition (Elmore et al. 2004, Hein et al. 2008). While male bats in our study did not select roosts with specific thermal characteristics, female bats seem to use less torpor and prefer warmer roosts than males while pregnant or raising young, and females typically aggregate in social maternity colonies rather than roosting solitarily (Hamilton and Barclay 1994, Kerth et al. 2001, Ruczyński 2006). Compared to males, then, roost selection by females will likely be governed more strongly by roost temperature (though social thermoregulation via huddling can influence temperatures within roosts more than a roost’s physical and environmental characteristics; Pretzlaff et al. 2010; Willis and Brigham, 2007). Further research on the roles of sex, age, and reproductive condition on torpor use in heterotherms (and thus habitat selection by heterotherms) is likely to reveal important context for our findings.

Climate warming increases energy expenditure for many animals, including both poikilotherms (Pörtner and Knust 2007, Dillon et al. 2010) and homeotherms (Humphries et al. 2002, Şekercioğlu et al. 2012). However, the degree to which climate warming will impact heterotherms is poorly understood, largely due to a lack of data on relationships between ambient temperature, torpor use, and thermolability that is needed to accurately model the influence of ambient temperature on heterotherm metabolism (Levesque et al. 2016). Our results indicate that temperature-dependent use of torpor may stabilize energy expenditure, and thus buffer against the energetic costs associated with variable ambient temperatures. However, most of the energetic savings from heterothermy arise during periods of cold. Increased temperatures due to climate change may thus reduce the relative energetic benefits of heterothermy compared to homeothermy, as homeotherms experience fewer and milder periods of cold.

In conclusion, we showed that a heterothermic bat selected neither warm nor cool roosts, because bats can modulate torpor use to stabilize energy expenditure over the course of a day. Unlike homeotherms, bats face little pressure to select warm habitats to avoid heat loss during periods of inactivity—when maintaining a high, stable body temperature becomes energetically costly, bats can calibrate the duration of torpor such that energy expenditure stays constant through a wide range of ambient temperature. Although such fine-tuning of torpor use to stabilize daily energy expenditure is intuitive, it has not been demonstrated in previous studies to the best of our knowledge.

## Acknowledgements

Many thanks to L. Boodoo, C. McFarland, E. Greene, B. Tabor, and B. Phillips for help with fieldwork; L. Shoemaker, D. Laughlin, J. Rick, and D. Goodhouse for helpful comments on pre-submission versions of this manuscript; and P. Ortegon, D. Licht, M. Wiles, D. Austin, B. Phillips, E. Thomas, and A. Stover for their logistical support of this project. Research funding was provided by the National Park Service, the Department of Zoology and Physiology at the University of Wyoming, the Berry Ecology Center, the American Society of Mammalogists, Prairie Biotic Research, Inc., and the Wyoming Chapter of The Wildlife Society. The findings and conclusions in this article are those of the authors and do not necessarily represent the views of the U.S. Fish and Wildlife Service or the National Park Service. We conducted field research on the traditional lands of the Lakȟóta, Sahnish, Tsitsistas, Hinono’eino, K’oigu, and Ná’ishą peoples. The Lakȟóta people know this land as Ȟe Sápa and Pahá Sápa, which was taken by the United States in the Agreement of 1877 in violation of the 1868 Fort Laramie Treaty.

## Data Availability

Data and code used in analyses for this paper will be archived on *Zenodo* upon acceptance of this manuscript and are available upon request by editors or reviewers.

## Author Contributions

JA, JG, and MD conceived and designed the study; JA, DK, and IA obtained funding for the study; JA collected and analyzed the data and led writing of the manuscript. All authors contributed to manuscript drafts and gave final approval for publication.

## Appendix S1: Supplementary Data

### Descriptive Statistics for Rock vs. Tree Roost Structures

During the day, rock crevices averaged 20.1°C (range: 16.5° – 24.2°C) while tree roost structures averaged 18.8°C (range: 16.1° – 25.5°C). Daily maximum temperatures within rock crevices averaged 26.1°C (range: 17.9° – 40.8°C), while daily maximum temperatures within tree roost structures averaged 28.3°C (range: 21.0° – 52.1°C). Temperatures within rock crevices peaked at 1441 hrs on average (range = 1005 – 1742 hrs), while temperatures within tree roost structures peaked at 1357 hrs on average (range = 1056 – 1659 hrs). Ambient temperature strongly influenced temperatures within roost structures. Temperatures within rock crevices at each hour (in °C) followed the equation 7.67 + 0.73*ambient temperature (*R*^2^ = 0.54), while temperatures within tree roost structures at each hour followed the equation 1.63 + 1.00*ambient temperature (*R*^2^ = 0.63).

Temperatures within used rock crevices averaged 20.5°C (range: 16.8° – 23.3°C) while temperatures within available rock crevices averaged 19.9°C (range: 16.5° – 24.2°C). Temperatures within used tree roosts averaged 18.6°C (range: 17.4° – 20.4°C) while temperatures within available tree cavities averaged 19.2°C (range: 16.1° – 25.5°C) and temperatures within available spaces under sloughing bark averaged 18.4°C (range: 16.1° – 21.0°C).

Temperatures within used rock crevices peaked on average at 1414 hrs (range: 1105 – 1719 hrs), while temperatures within available rock crevices peaked on average at 1458 hrs (range: 1005 – 1742 hrs). Temperatures within used tree roosts peaked on average at 1447 hrs (range: 1125 – 1659 hrs), while temperatures within available tree cavities peaked on average at 1410 hrs (range: 1120 – 1608 hrs) and temperatures within available spaces under sloughing bark peaked on average at 1349 hrs (range: 1056 – 1608 hrs).

The standard deviation of temperatures within used rock crevices was 6.7°C (range: 4.3° – 10.0°C), while the standard deviation of temperatures within available rock crevices was 6.2°C (range: 3.2° - 11.0°C). The standard deviation of temperatures within used tree roosts was 7.7°C (range: 6.7° - 9.1°C), while the standard deviation of temperatures within available tree cavities was 8.7°C (range: 5.9° - 16.4°C) and within available spaces under sloughing bark was 7.7°C (range: 6.5° - 11.0°C).

There was no difference in ambient temperature between days where rock crevices were used and days where tree roost structures were used (Mann-Whitney U = 299; *p* = 0.968).

---

**Table S1.** Information on torpor use by bats tracked during our study, including an ID number for each individual, the dates for which we have data, the mass of bats at time of capture, the timing of torpor entry and exit for morning and afternoon bouts of torpor, the duration of periods of periods of torpor in both mornings and afternoons, and the total duration of torpor across the day.

| Bat ID | Date | Mass<br>(grams) | AM Torpor<br>Start Time<br>(hrs) | AM Torpor<br>End Time<br>(hrs) | Duration of<br>AM Torpor<br>(mins) | PM Torpor<br>Start Time<br>(hrs) | PM Torpor<br>End Time<br>(hrs) | Duration of<br>PM Torpor<br>(mins) | Total Torpor<br>Duration<br>(mins) |
| --- | --- | --- | --- | --- | --- | --- | --- | --- | --- |
| 172_063 | 8/5/2017 | 6.02 | 517 | 1456 | 579 | 2013 | 2055 | 42 | 621 |
| 172_063 | 8/6/2017 | 6.02 | 451 | 1210 | 439 | 1910 | 2037 | 87 | 526 |
| 172_063 | 8/7/2017 | 6.02 | 2245 | 1557 | 1032 | 1840 | 2044 | 124 | 1156 |
| 172_904 | 6/28/2018 | 6.75 | 425 | 733 | 188 | 1825 | 2057 | 125 | 313 |
| 172_904 | 6/29/2018 | 6.75 | 419 | 1037 | 378 | 1603 | 2114 | 277 | 655 |
| 172_904 | 7/3/2018 | 6.75 | 525 | 944 | 259 | 1834 | 2029 | 115 | 374 |
| 172_904 | 7/4/2018 | 6.75 | 412 | 1446 | 634 | 1709 | 2122 | 253 | 887 |
| 172_904 | 7/5/2018 | 6.75 | 424 | 1458 | 597 | 1930 | 2043 | 73 | 670 |
| 172_904 | 7/6/2018 | 6.75 | 511 | 1016 | 305 | - | - | 0 | 305 |
| 172_904 | 7/7/2018 | 6.75 | 438 | 818 | 220 | - | - | 0 | 220 |
| 172_692 | 7/13/2018 | 6.92 | 445 | 830 | 225 | 1936 | 2043 | 67 | 292 |
| 172_692 | 7/14/2018 | 6.92 | 435 | 815 | 220 | - | - | 0 | 220 |
| 172_632 | 7/20/2018 | 8.04 | 426 | 1102 | 396 | 1916 | 2041 | 85 | 481 |
| 172_753 | 7/27/2018 | 8.16 | 133 | 2045 | 1152 | - | - | 0 | 1152 |
| 172_753 | 7/28/2018 | 8.16 | 2300 | 2031 | 1291 | - | - | 0 | 1291 |
| 172_453 | 8/4/2018 | 7.1 | 449 | 959 | 310 | 1915 | 2039 | 84 | 394 |
| 172_784 | 8/4/2018 | 7.53 | 442 | 1028 | 346 | 1951 | 2023 | 32 | 378 |
| 172_453 | 8/5/2018 | 7.1 | 459 | 1156 | 417 | 1613 | 2028 | 255 | 672 |
| 172_784 | 8/5/2018 | 7.53 | 445 | 1100 | 375 | 1852 | 2019 | 87 | 462 |
| 172_453 | 8/6/2018 | 7.1 | 441 | 916 | 275 | 1823 | 2034 | 131 | 406 |
| 172_784 | 8/6/2018 | 7.53 | 449 | 1003 | 314 | - | - | 0 | 314 |
| 172_453 | 8/7/2018 | 7.1 | 444 | 1041 | 357 | - | - | 0 | 357 |
| 172_784 | 8/7/2018 | 7.53 | 502 | 850 | 228 | - | - | 0 | 228 |
| 172_063 | 8/8/2018 | 6.02 | 2335 | 1427 | 892 | 1737 | 2009 | 152 | 1044 |
| 172_453 | 8/8/2018 | 7.1 | 451 | 839 | 228 | - | - | 0 | 228 |
| 172_784 | 8/8/2018 | 7.53 | 439 | 852 | 253 | - | - | 0 | 253 |
| 172_453 | 8/10/2018 | 7.1 | 456 | 843 | 227 | - | - | 0 | 227 |

**Fig. S1.**
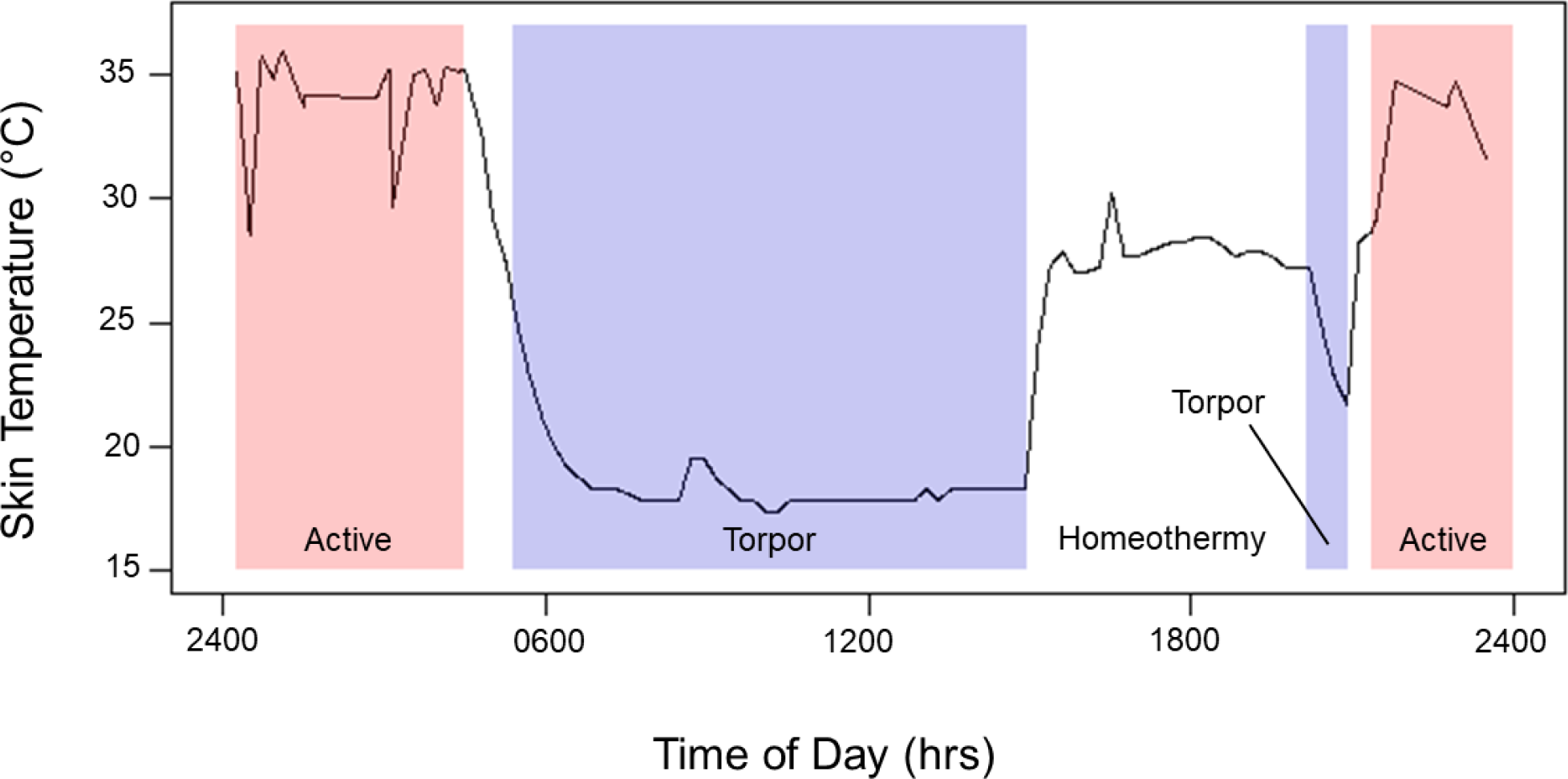
An example of raw skin temperature data that we used to delineate bouts of torpor. Periods of time in red blocks represent periods of activity (flying, foraging, etc.), periods of time in blue blocks represent periods of torpor, and periods in white represent periods of homeothermy or transition between torpor and homeothermy/activity. To delineate bouts of torpor, we used the definition suggested in Barclay et al. (2001).

**Fig. S2.**
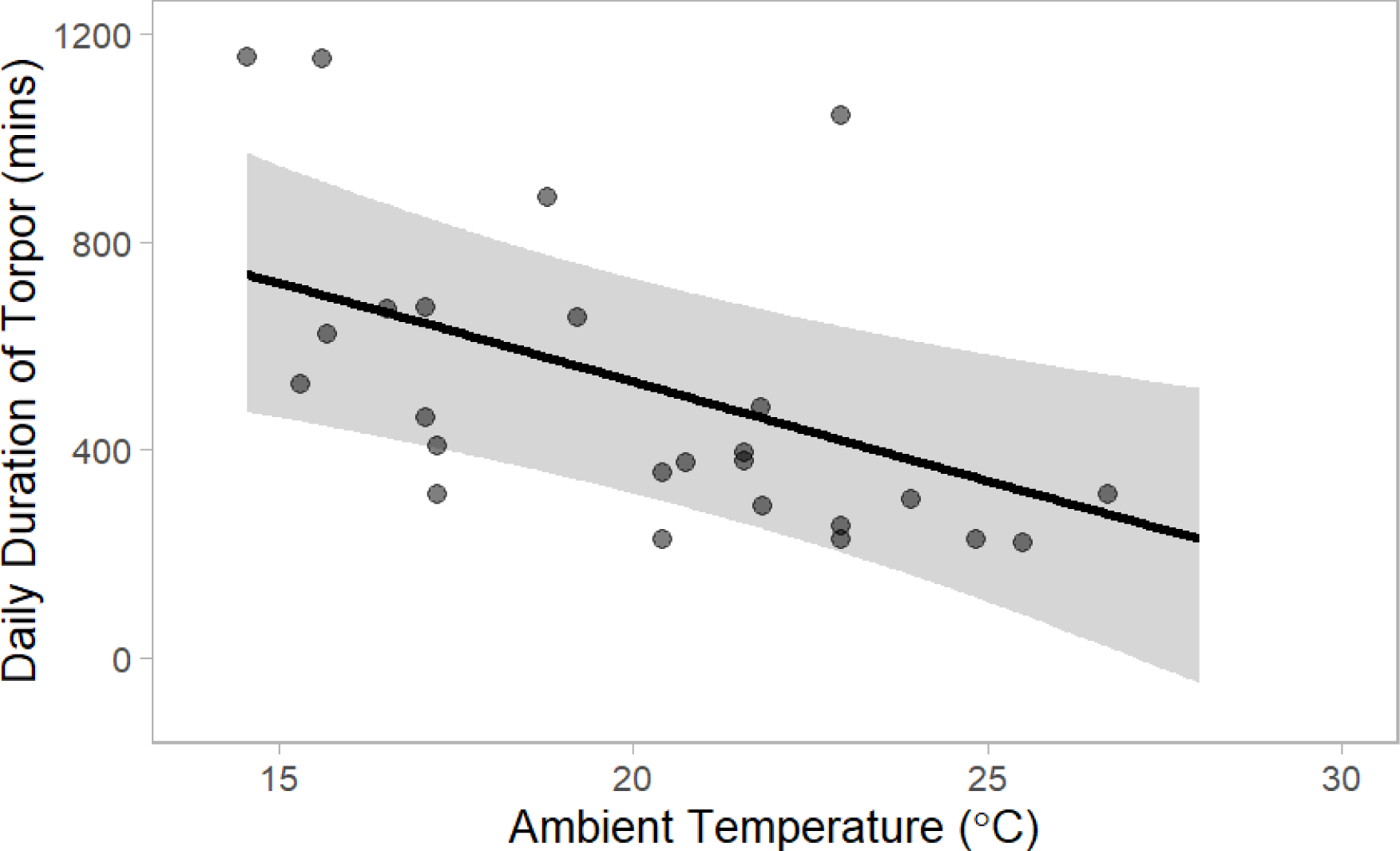
Scatter plot illustrating the conditional effect of daily mean ambient temperature on the total duration of bouts of torpor during the day. Each point is based on observed data and represents one day. The line represents the regression line for this relationship and the grey band represents 95% credible intervals around this line. Credible intervals for this conditional effect did not cross zero (parameter estimate: −37.4 min; 95% credible intervals: −64.0 – −12.6 min), indicating that bats spent ca. 37 minutes less in torpor per day for each additional 1°C in daily mean ambient temperature between 0445 hrs and 2100 hrs.

